## Extended Data for "Optimizing a Human Monoclonal Antibody for Better Neutralization of SARS-CoV-2"

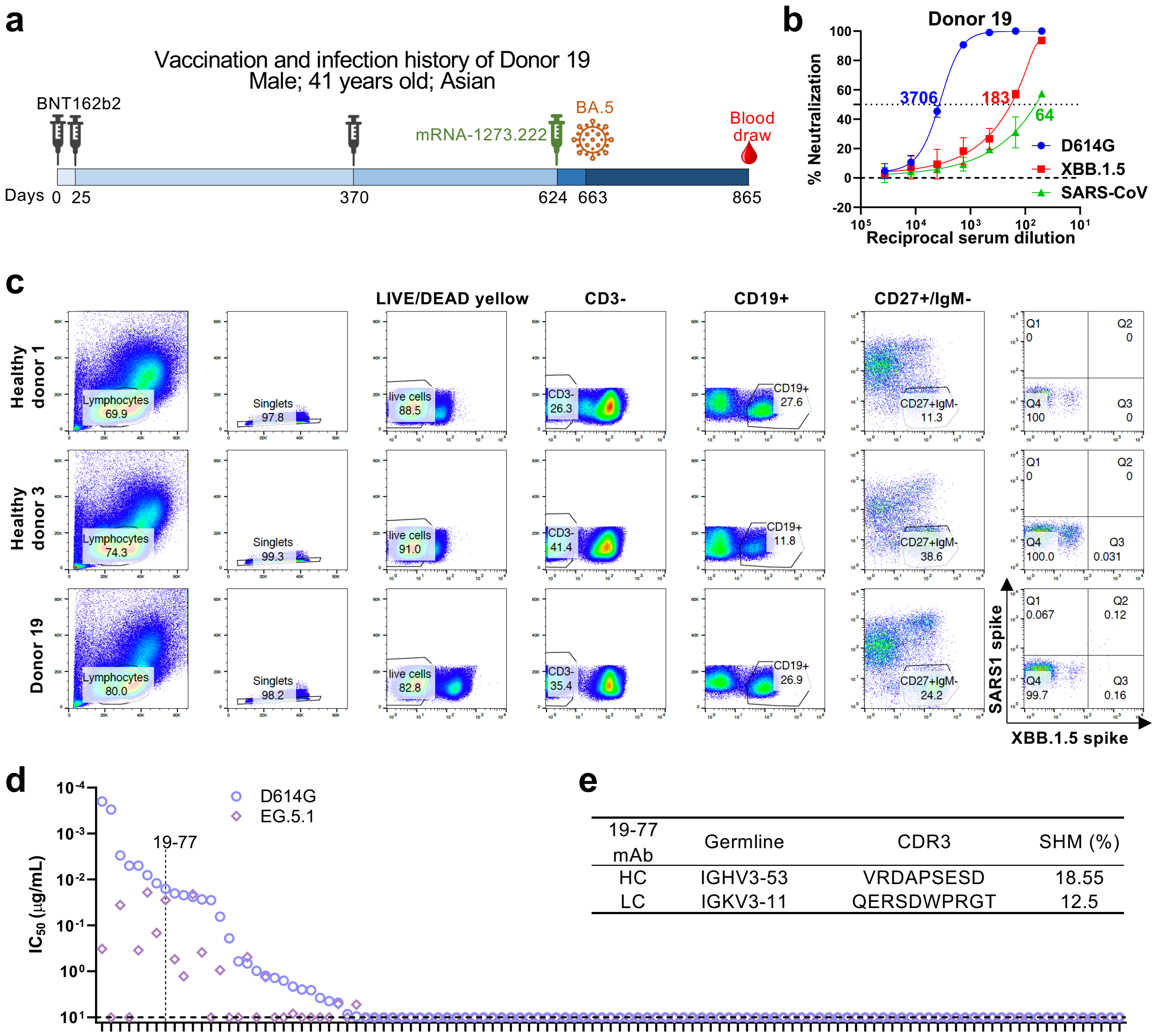


| 19-77 mAb | Germline | CDR3 | SHM (%) |
|---|---|---|---|
| HC | IGHV3-53 | VRDAPSESD | 18.55 |
| LC | IGKV3-11 | QERSDWPRGT | 12.5 |

**Extended Data Fig. 1 | Clinical information and sorting strategy.**

**a.** Clinical information of donor 19, whose blood sample was collected for neutralizing antibody evaluation and B cell sorting.

**b.** Neutralization activity of serum from donor 19 against D614G, XBB.1.5, and SARS-CoV. Neutralization $ID_{50}$ titer against each virus is denoted. Data are shown as mean ± SEM (standard error of the mean) from technical triplicates.

**c.** Sorting strategy used to isolate XBB.1.5 spike and/or SARS-CoV spike specific B cells. B cells from Q2 and Q3 were collected and applied for downstream 10X Genomics analysis. Numbers in gates represent cell percentages.

**d.** Neutralizing $IC_{50}$ values of the antibodies from donor 19 against D614G and EG.5.1. 19-77 is highlighted with a dotted line.

**e.** Germline genes, CDR3 amino acid sequences, and SHM percentages of 19-77 heavy and light chains.

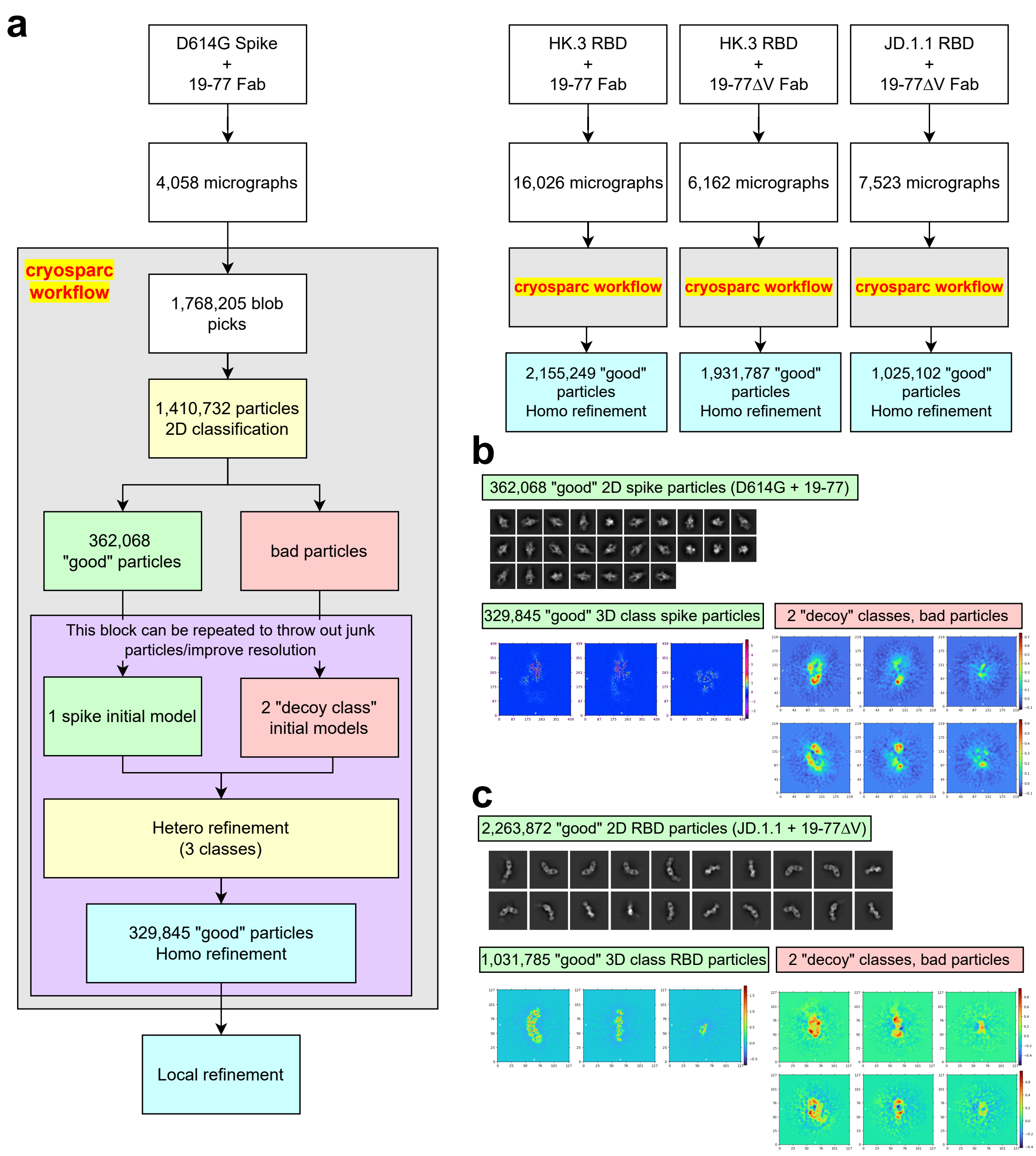


**Extended Data Fig. 2 | Cryo-EM processing workflow and sample class averages**

**a.** Cryo-EM single particle processing workflow for spike+antibody and RBD+antibody complexes. A standard cryo-EM single-particle processing pipeline was applied individually to each dataset of spike+antibody and RBD+antibody complexes.

**b.** Representative 2D and 3D classes of spike+antibody particles are shown using D614G and 19-77 as example.

**c.** Representative 2D and 3D classes of RBD+antibody particles are shown using JD.1.1 RBD and 19-77ΔV as example.

a

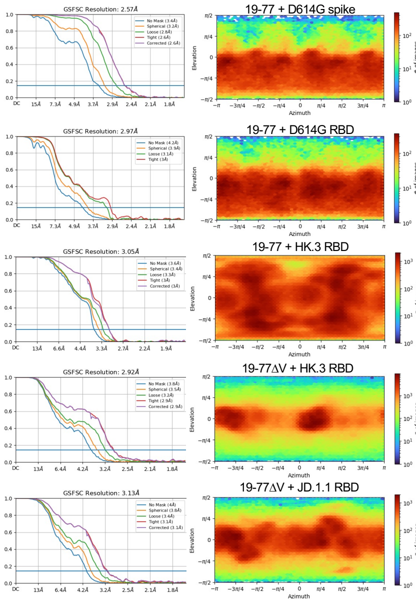


b

**Cryo-EM data collection, refinement, and validation statistics for 19-77 samples**

| | | | | |
|---|---|---|---|---|
| Antibody | 19-77 | 19-77 | 19-77ΔV | 19-77ΔV |
| Antigen | D614G RBD | HK.3 RBD | HK.3 RBD | JD.1.1 RBD |
| EMBD | EMD-45543 | EMD-45544 | EMD-45545 | EMD-45546 |
| PDB | 9CFE | 9CFF | 9CFG | 9CFH |
| **Data collection and processing** | | | | |
| Magnification | 105,000 | 105,000 | 105,000 | 105,000 |
| Voltage (kV) | 300 | 300 | 300 | 300 |
| Electron exposure (e-/Å²) | 58.06 | 58.06 | 58.06 | 58.06 |
| Defocus rang (μm) | -0.8 to -2.0 | -0.8 to -2.0 | -0.8 to -2.0 | -0.8 to -2.0 |
| Pixel size (Å/pix) | 0.83 | 0.83 | 0.83 | 0.83 |
| Symmetry impose | C1 | C1 | C1 | C1 |
| Final particle images (no.) | 329,845 | 2,155,249 | 1,931,787 | 1,025,102 |
| Map resolution | 3.1Å | 3.3Å | 3.2Å | 3.4Å |
| FSC threshold (loose mask) | 0.143 | 0.143 | 0.143 | 0.143 |
| **Refinement** | | | | |
| Model composition | | | | |
| Non-hydrogen atoms | 6382 | 6400 | 6402 | 6394 |
| Protein residues | 416 | 417 | 417 | 417 |
| Ligands | | | | |
| R.m.s. deviations | | | | |
| Bond lengths (Å) | 0.013 (2) | 0.004 (0) | 0.004 (0) | 0.004 (0) |
| Bond angles (˚) | 1.928 (31) | 0.625 (3) | 0.744 (0) | 0.747 (3) |
| Validation | | | | |
| MolProbity score | 1.32 | 1.74 | 1.47 | 1.25 |
| Clashscore | 0.31 | 4.68 | 5.63 | 2.50 |
| Poor rotamers (%) | 1.98 | 1.69 | 0.28 | 0.00 |
| Ramachandran plot | | | | |
| Favored (%) | 92.20 | 95.38 | 97.08 | 96.59 |
| Allowed (%) | 6.83 | 4.38 | 2.92 | 3.16 |
| Disallowed (%) | 0.98 | 0.24 | 0.0 | 0.24 |
| B factors (Å) | | | | |
| Protein | 41.52 | 65.72 | 147.56 | 58.15 |

c

**X-ray diffraction data collection and refinement statistics**

| | |
|---|---|
| Antibody | 19-77 Fab |
| Antigen | EG.5.1 RBD |
| PDB ID | 9CAV |
| **Data Collection** | |
| Space group | $P2_1$ |
| Cell constants | 94.9 116.2 109.2 90 110.4 90 |
| Resolution range (Å) | 35.2 - 3.2 (3.1- 3.0)* |
| Total reflections | 86848 (514) |
| Unique reflections | 43474 (1270) |
| Completeness (%) | 97.74 (94.91) |
| Redundancy | 2.0 (2.0) |
| $I/\sigma$ | 3.22 (1.29) |
| R-merge | 0.085 (0.67) |
| CC1/2 | 0.69 (0.40) |
| **Refinement** | |
| Resolution range (Å) | 34.5 - 3.2 |
| Number of complexes per asymmetric unit | 2 |
| Rwork/Rfree | 0.25/0.29 |
| Number of atoms | |
| Protein | 8940 |
| Ligands | 0 |
| Water | 0 |
| B-factors (Å²) | |
| Protein | 36.0 |
| Ligands | - |
| Water | - |
| R.M.S deviations | |
| Bond lengths (Å) | 0.012 |
| Bond angles (˚) | 1.64 |
| Ramachandran statistics | |
| Favored (%) | 93.85 |
| Allowed (%) | 6.15 |
| Outliers (%) | 0 |

d

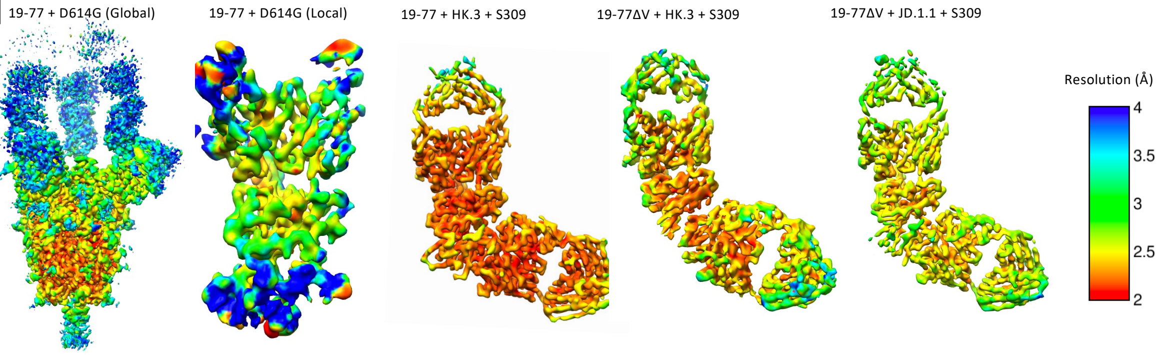


e

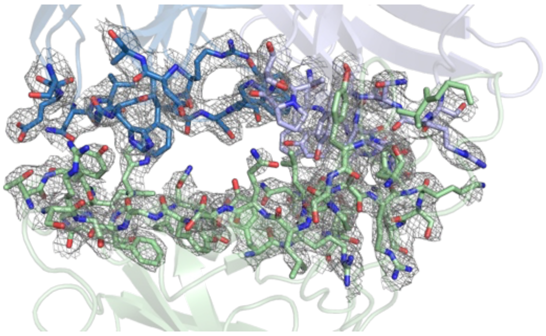

**Extended Data Fig. 3 | Cryo-EM and X-ray data for mAbs 19-77 and 19-77ΔV in complex with SARS-CoV-2 spike trimers or RBDs.**

**a.** Global refinement Fourier Shell Correction curves showing the overall resolution of the indicated complexes.

**b.** Cryo-EM data collection and model refinement of the indicated complexes.

**c.** X-ray diffraction data collection and refinement statistics.

**d.** Global and local resolution of Cryo-EM structures.

**e.** Composite omit map showing the electron density at the interface between EG5.1 RBD and 19-77 Fab. The RBD is depicted in pale green, with the Fab light and heavy chains shown in different shades of blue. The map is contoured at 1.5σ.

**a**

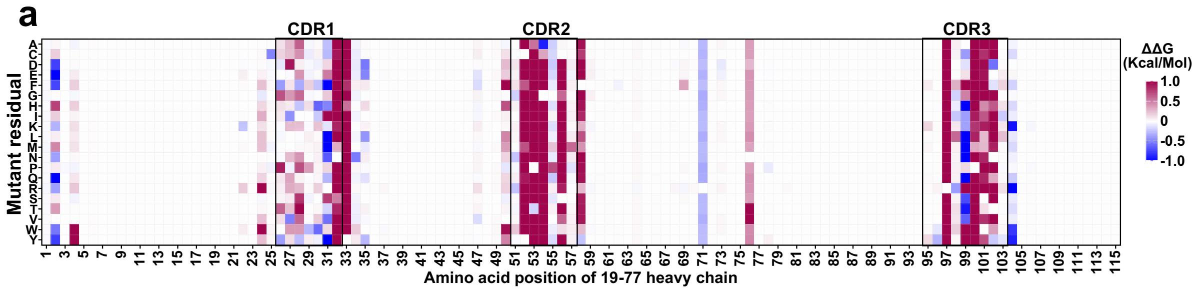


**b**

19-77 HC

| IC50 (μg/mL) | JD.1.1 | HK.3 | EG.5 | EG.5-A475V |
|---|---|---|---|---|
| 19-77 | >10 | 0.229 | 0.098 | >10 |
| E1A | >10 | 0.450 | 0.137 | >10 |
| V2A | >10 | 0.202 | 0.060 | >10 |
| S25A | >10 | 0.178 | 0.088 | >10 |
| E26A | >10 | 0.570 | 0.281 | >10 |
| L27A | >10 | 2.309 | 0.523 | >10 |
| I28A | >10 | 0.940 | 0.156 | >10 |
| S30A | >10 | 0.114 | 0.046 | >10 |
| R31A | >10 | 0.210 | 0.069 | >10 |
| N32A | >10 | >10 | >10 | >10 |
| Y33A | >10 | >10 | >10 | >10 |
| Y52A | >10 | 0.463 | 0.030 | >10 |
| P53A | >10 | 0.887 | 0.046 | >10 |
| G54A | >10 | 2.377 | 0.923 | >10 |
| S56A | >10 | 3.750 | 1.092 | >10 |
| S57A | >10 | 0.068 | 0.059 | >10 |
| F58A | >10 | >10 | 2.935 | >10 |
| R71A | 0.169 | 0.009 | 0.009 | 0.176 |
| N76A | >10 | 0.286 | 0.049 | >10 |
| R97A | >10 | >10 | >10 | >10 |
| D98A | >10 | 2.840 | 0.550 | >10 |
| P100A | >10 | 0.241 | 0.584 | >10 |
| S101A | >10 | 0.168 | 0.071 | >10 |
| E102A | >10 | 0.619 | 0.062 | >10 |

**c**

19-77 LC

| IC50 (μg/mL) | JD.1.1 | HK.3 | EG.5 | EG.5-A475V |
|---|---|---|---|---|
| 19-77 | >10 | 0.229 | 0.098 | >10 |
| I2A | >10 | 0.994 | 0.121 | >10 |
| Q27A | >10 | 0.280 | 0.081 | >10 |
| N28A | >10 | >10 | >10 | >10 |
| I29A | >10 | 0.578 | 0.036 | >10 |
| G30A | >10 | 0.294 | 0.066 | >10 |
| H31A | >10 | >10 | 1.244 | >10 |
| F32A | >10 | >10 | 0.440 | >10 |
| D50A | >10 | 2.456 | 0.225 | >10 |
| S67A | >10 | 0.528 | 0.127 | >10 |
| E68A | >10 | 1.724 | 0.315 | >10 |
| S92A | >10 | 0.135 | 0.074 | >10 |
| D93A | >10 | 0.388 | 0.098 | >10 |
| W94A | >10 | >10 | >10 | >10 |
| P95A | >10 | >10 | 0.469 | >10 |

**d**

| IC50 (μg/mL) | JD.1.1 | HK.3 | EG.5 | EG.5-A475V |
|---|---|---|---|---|
| 19-77 | >10 | 0.170 | 0.062 | >10 |
| R71A | 0.065 | 0.008 | 0.006 | 0.337 |
| R71C | 0.606 | 0.014 | 0.008 | 0.336 |
| R71D | 0.420 | 0.010 | 0.011 | 1.211 |
| R71E | 0.472 | 0.009 | 0.010 | 0.333 |
| R71F | 0.469 | 0.007 | 0.006 | 0.168 |
| R71G | 0.546 | 0.005 | 0.006 | 0.850 |
| R71H | 0.310 | 0.007 | 0.008 | 0.301 |
| R71I | 0.285 | 0.010 | 0.011 | 0.206 |
| R71K | >10 | 0.048 | 0.030 | 3.394 |
| R71L | 0.095 | 0.008 | 0.013 | 0.115 |
| R71M | 0.241 | 0.008 | 0.012 | 0.195 |
| R71N | 0.324 | 0.010 | 0.006 | 0.321 |
| R71P | 0.259 | 0.010 | 0.007 | 0.82 |
| R71Q | 0.245 | 0.003 | 0.012 | 0.214 |
| R71S | 0.319 | 0.008 | 0.015 | 0.237 |
| R71T | 0.251 | 0.007 | 0.015 | 0.155 |
| R71V | 0.095 | 0.004 | 0.015 | 0.06 |
| R71W | 0.615 | 0.006 | 0.019 | 1.054 |
| R71Y | 0.434 | 0.005 | 0.006 | 0.234 |

| <0.01 | 0.01-0.1 | 0.1-1 | 1-10 | >10 |
|---|---|---|---|---|

**Extended Data Fig. 4 | Neutralizing $IC_{50}$ values of 19-77 mutants.**

**a.** Free energy change (ΔΔG) for saturation mutagenesis of the 19-77 heavy chain. Blue represents beneficial mutations, while red indicates mutations with adverse effects.

**b.** Neutralization $IC_{50}$ values of 19-77 carrying the indicated mutations in the heavy chain.

**c.** Neutralization $IC_{50}$ values of 19-77 carrying the indicated mutations in the light chain.

**d.** Neutralization $IC_{50}$ values of 19-77 carrying various amino acid substitutions at R71 in the heavy chain.

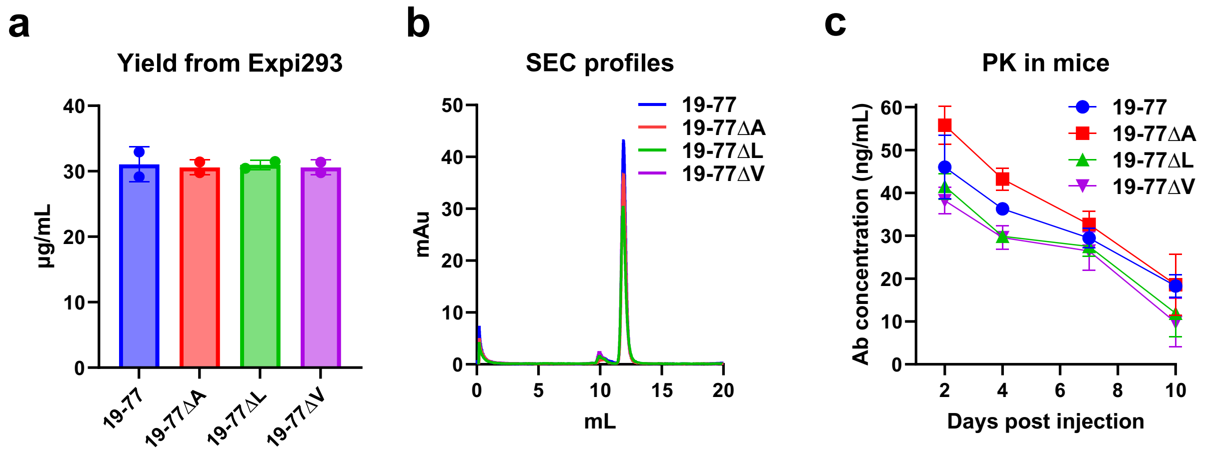


**Extended Data Fig. 5 | Biological properties of 19-77.**

- **a.** Yields of 19-77 and 19-77ΔA/L/V from transiently transfected Expi293 cells at day 4 post-transfection, data show from technical duplicates.
- **b.** Size exclusion chromatography (SEC) profiles of 19-77 and 19-77ΔA/L/V antibodies.
- **c.** Pharmacokinetics of 19-77 and 19-77ΔA/L/V in mice over 10 days after intraperitoneal injection. Data are shown as mean ± SEM (standard error of the mean) from technical triplicates.

| Abs | | | | Ab IC50 (ug/mL) | | | | | |
|---|---|---|---|---|---|---|---|---|---|
| | | | | D614G | EG.5.1 | BA.2.86 | JN.1 | HK.3 | JD.1.1 |
| VH3-53 | BD56-1302 | RBD class 1 | WT | 0.014 | 0.014 | 0.007 | 0.186 | 0.202 | >10 |
| | | | R71A | 0.011 | 0.012 | 0.005 | 0.036 | 0.060 | >10 |
| | | | R71L | 0.012 | 0.019 | 0.006 | 0.073 | 0.152 | >10 |
| | | | R71V | 0.009 | 0.016 | 0.005 | 0.038 | 0.115 | >10 |
| | BD56-1854 | RBD class 1 | WT | 0.008 | 0.027 | 0.002 | 0.022 | 0.111 | 10 |
| | | | R71A | 0.006 | 0.004 | 0.001 | 0.004 | 0.031 | 0.189 |
| | | | R71L | 0.004 | 0.008 | 0.002 | 0.011 | 0.053 | 0.697 |
| | | | R71V | 0.005 | 0.005 | 0.001 | 0.005 | 0.048 | 0.977 |
| | Omi3 | RBD class 1 | WT | 0.011 | >10 | 0.467 | >10 | >10 | >10 |
| | | | R71A | 0.009 | 0.668 | 0.021 | 0.685 | 1.505 | >10 |
| | | | R71L | 0.009 | 0.991 | 0.048 | 1.324 | 2.232 | >10 |
| | | | R71V | 0.005 | 0.172 | 0.020 | 0.331 | 0.912 | >10 |
| | 19-79 | RBD class 1 | WT | 0.013 | 0.073 | 0.007 | >10 | >10 | >10 |
| | | | R71A | 0.011 | 0.042 | 0.002 | 0.116 | 0.165 | >10 |
| | | | R71L | 0.016 | 0.160 | 0.006 | 0.484 | 0.582 | >10 |
| | | | R71V | 0.011 | 0.062 | 0.003 | 0.124 | 0.162 | >10 |
| VH3-66 | BD57-0129 | RBD class 1 | WT | 0.013 | 0.003 | 0.003 | >10 | >10 | >10 |
| | | | R71A | 0.013 | 0.003 | 0.002 | 0.028 | 0.107 | >10 |
| | | | R71L | 0.015 | 0.005 | 0.003 | 0.064 | 0.132 | >10 |
| | | | R71V | 0.013 | 0.003 | 0.002 | 0.035 | 0.090 | >10 |
| | BD515 | RBD class 1 | WT | 0.022 | >10 | 0.166 | >10 | >10 | >10 |
| | | | R71A | 0.017 | >10 | 0.115 | 0.322 | >10 | >10 |
| | | | R71L | 0.043 | >10 | 0.157 | 0.913 | >10 | >10 |
| | | | R71V | 0.016 | 0.955 | 0.028 | 0.098 | 1.818 | >10 |
| VH3-74 | C68.59 | SD1 | WT | 0.048 | 0.108 | >10 | >10 | 0.081 | 0.073 |
| | | | R71A | 0.021 | 0.173 | >10 | >10 | 0.105 | 0.099 |
| | | | R71V | 0.025 | 0.179 | >10 | >10 | 0.085 | 0.120 |
| | | | R71L | 0.083 | 0.341 | >10 | >10 | 0.149 | 0.150 |
| VH3-9 | Omi42 | RBD class 1 | WT | 0.020 | >10 | 0.021 | 0.066 | >10 | >10 |
| | | | R71A | 0.056 | >10 | >10 | >10 | >10 | >10 |
| | | | R71V | 0.048 | >10 | >10 | >10 | >10 | >10 |
| | | | R71L | 0.057 | >10 | >10 | >10 | >10 | >10 |

| <0.01 | 0.01-0.1 | 0.1-1 | 1-10 | >10 |
|---|---|---|---|---|

**Extended Data Fig. 6 | R71/A/L/V mutations rescue the neutralizing activities of VH3-53 and VH3-66 antibodies against SARS-CoV-2 escaping variants.** The figure shows neutralization $IC_{50}$ values of the indicated antibodies and their R71A/L/V mutants.

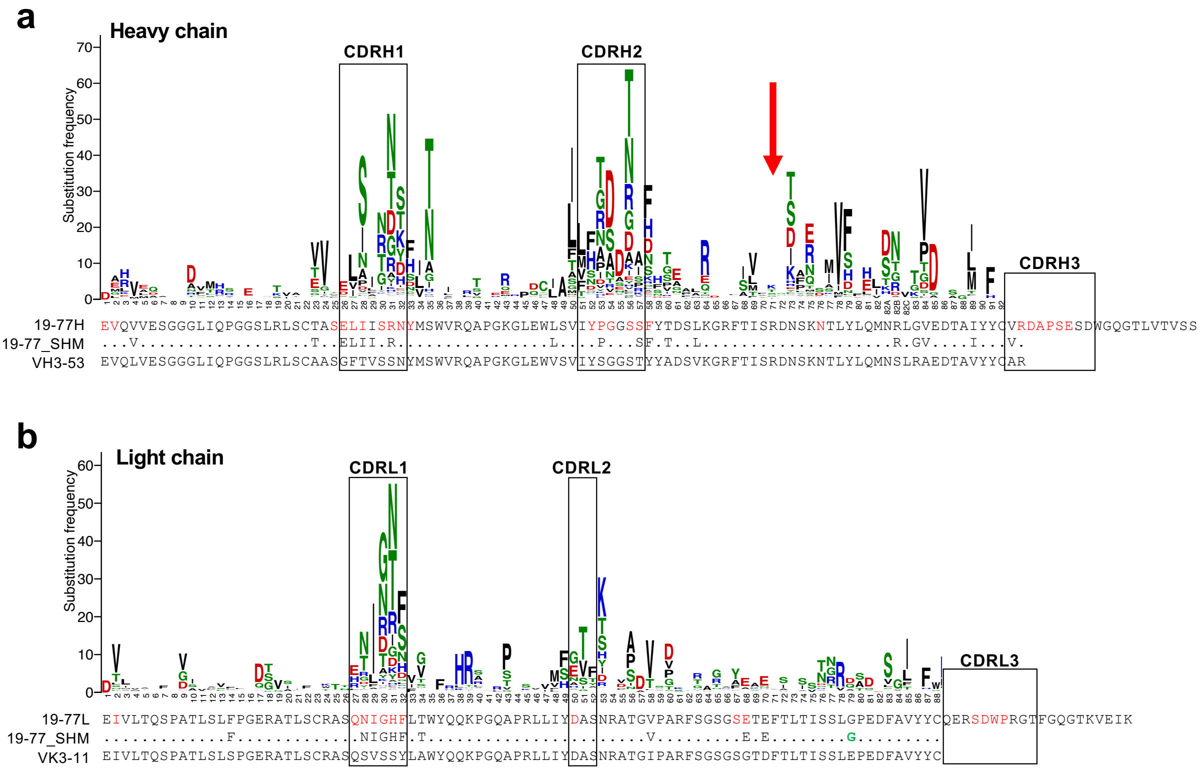


**Extended Data Fig. 7 | Gene-specific substitution profile for mAb 19-77.**

**a.** The gene-specific substitution profiles (GSSP) for IGHV3-53 and the SHMs of 19-77 heavy chain. The red residues indicate the paratope residues of 19-77 heavy chain. R71 is highlighted by a red arrow.

**b.** GSSP for IGKV3-11 and the SHMs of 19-77 heavy chain. The red residues indicate the paratope residues of 19-77 light chain. The green residue indicates a rare mutation in 19-77 light chain.

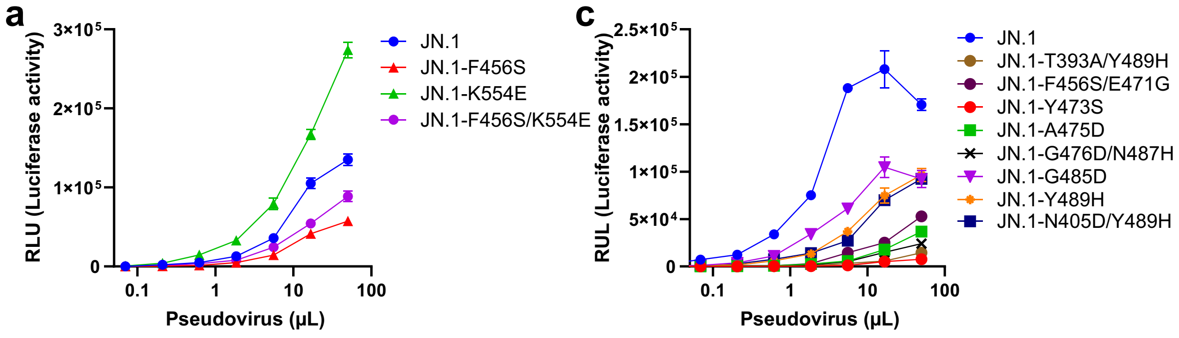


**b**

| Neutralization ($IC_{50}$, μg/mL) | 19-77 | 19-77ΔV | hACE2 |
|---|---|---|---|
| JN.1 | 0.101 | 0.013 | 0.315 |
| JN.1-E554K | 0.104 | 0.009 | 0.167 |
| JN.1-F456S | >10 | 0.675 | 0.304 |
| JN.1-F456S/K554E | >10 | 0.424 | 0.695 |

| <0.01 | 0.01-0.1 | 0.1-1 | 1-10 | >10 |
|---|---|---|---|---|

**d**

| Neutralization ($IC_{50}$, μg/mL) | 19-77 | 19-77ΔA | 19-77ΔL | 19-77ΔV | hACE2 |
|---|---|---|---|---|---|
| JN.1 | 0.047 | 0.008 | 0.009 | 0.008 | 0.088 |
| JN.1-T393A/Y489H | >10 | >10 | >10 | >10 | >10 |
| JN.1-F456S/E471G | >10 | >10 | >10 | >10 | 9.039 |
| JN.1-Y473S | >10 | >10 | >10 | >10 | >10 |
| JN.1-A475D | >10 | >10 | >10 | >10 | >10 |
| JN.1-G476D/N487H | >10 | >10 | >10 | >10 | >10 |
| JN.1-G485D | >10 | 0.074 | 0.113 | 0.084 | 1.094 |
| JN.1-Y489H | >10 | 0.363 | 0.519 | 0.976 | 2.345 |
| JN.1-N405D/Y489H | >10 | >10 | >10 | >10 | 2.967 |

**Extended Data Fig. 8 | Infectivity and neutralization activity of 19-77ΔV escape variants in the context of VSV pseudotyped viruses.**

**a.** Infectivity of the indicated pseudotyped escape variants selected by 19-77ΔV from authentic JN.1 in Vero-E6-TMPRSS2-T2A-ACE2 cells. Data are shown as mean ± SEM (standard error of the mean) from technical triplicates.

**b.** Infectivity of the indicated pseudotyped escape variants selected by 19-77ΔV from replication-competent VSV-JN.1 in Vero-E6-TMPRSS2-T2A-ACE2 cells. Data are shown as mean ± SEM from technical triplicates.

**c.** Neutralization $IC_{50}$ values of 19-77, 19-77ΔV, and hACE2 against the indicated pseudotyped escape variants.

**d.** Neutralization $IC_{50}$ values of 19-77, 19-77ΔA/L/V, and hACE2 against the indicated pseudotyped escape variants.
